# The connectional anatomy of visual mental imagery: evidence from a patient with left occipito-temporal damage

**DOI:** 10.1101/2022.02.15.480510

**Authors:** Dounia Hajhajate, Brigitte Kaufmann, Jianghao Liu, Katarzyna Siuda-Krzywicka, Paolo Bartolomeo

**Affiliations:** Sorbonne Université, Institut du Cerveau - Paris Brain Institute - ICM, Inserm, CNRS, AP-HP, Hôpital de la Pitié-Salpêtrière, F-75013 Paris, France; Dassault Systèmes, France

**Keywords:** Perception & imagery, patients, cerebrovascular, behavioral, lesion mapping, white matter tractography

## Abstract

Most of us can use our “mind’s eye” to mentally visualize things that are not in our direct line of sight, an ability known as visual mental imagery. Extensive left temporal damage can impair patients’ visual mental imagery experience, but the critical locus of lesion is unknown. Our recent meta-analysis of 27 fMRI studies of visual mental imagery highlighted a well-delimited region in the left lateral midfusiform gyrus, which was consistently activated during visual mental imagery, and which we called the Fusiform Imagery Node (FIN). Here we describe the connectional anatomy of FIN in neurotypical participants and in RDS, a right-handed patient with an extensive occipitotemporal stroke in the left hemisphere. The stroke provoked right homonymous hemianopia, alexia without agraphia, and color anomia. Despite these deficits, RDS had normal subjective experience of visual mental imagery and reasonably preserved behavioral performance on tests of visual mental imagery of object shape, object color, letters, faces, and spatial relationships. We found that the FIN was spared by the lesion. We then assessed the connectional anatomy of the FIN in the MNI space and in the patient’s native space, by visualizing the fibers of the inferior longitudinal fasciculus (ILF) and of the arcuate fasciculus (AF) passing through the FIN. In both spaces, the ILF connected the FIN with the anterior temporal lobe, and the AF linked it with frontal regions. Our evidence is consistent with the hypothesis that the FIN is a node of a brain network dedicated to voluntary visual mental imagery. The FIN could act as a bridge between visual information and semantic knowledge processed in the anterior temporal lobe and in the language circuits.

## Introduction

Visual mental imagery denotes our ability to use our “mind’s eye” to mentally visualize things that are not in our direct line of sight. Brain-damaged patients with extensive left temporal damage often have impaired visual mental imagery (Bartolomeo, 2002, 2008; Bartolomeo, Hajhajate, Liu, & Spagna, 2020; Spagna, 2022), but the crucial lesion site in the temporal cortex is unknown. Our recent meta-analysis of 27 fMRI studies of visual mental imagery (Spagna, Hajhajate, Liu, & Bartolomeo, 2021) highlighted the importance of a well-delimited region within the FG4 field (Lorenz et al., 2015) of the left midfusiform gyrus, that we labeled Fusiform Imagery Node (FIN). This finding is consistent with the available neuroanatomical evidence from lesion neuropsychology, neuroimaging, and direct cortical stimulation, which indicates the left inferior temporal lobe as the region most commonly implicated in voluntary generated mental images (Liu, Spagna, & Bartolomeo, 2021). The localization of the FIN in the left ventral temporal cortex suggests a possible role as a bridge between domain-preferring visual regions (Mahon & Caramazza, 2011) and amodal semantic networks (Fairhall & Caramazza, 2013; Lambon Ralph, Jefferies, Patterson, & Rogers, 2017), perhaps including the language circuits (Bouhali et al., 2014).

However, neuroimaging evidence such as that coming from the Spagna et al’s (2021) meta-analysis is correlative, not causal. To establish a causal role of FIN in visual mental imagery, the study of brain-damaged patients is mandatory. Here we describe patient RDS, a 58-year-old, right-handed patient who 7 years before testing had an extensive left occipitotemporal stroke. The stroke provoked right homonymous hemianopia, alexia without agraphia, and color anomia. These deficits and their neuroanatomical correlates were extensively described in previous papers (Siuda-Krzywicka et al., 2019; Siuda-Krzywicka et al., 2020), and were stable at the time of the present testing. Despite his deficits, RDS had preserved mental imagery introspection and behavioral performance. To estimate the importance of FIN for visual mental imagery in this patient, we mapped it on the native space of his brain. Moreover, we assessed its connectional anatomy by mapping it and the lesion in the MNI and native spaces; we then visualized the fibers of the two most important long-range white matter tracts passing through the FIN, the inferior longitudinal fasciculus (ILF) and the arcuate fasciculus (AF). We chose to examine these two major pathways because of their likely role in linking the FIN to the processing of semantic information in the anterior temporal lobe (Lambon Ralph et al., 2017) and in the perisylvian language network (Bouhali et al., 2014).

## Methods

We used a French version (Santarpia et al., 2008) of the VVIQ questionnaire (Marks, 1973) to assess RDS’s subjective vividness of visual mental imagery. In Santarpia et al’s (2008) classification based on neurotypical controls, scores ≤46 indicate low imagery vividness, scores 47–65 indicate average vividness, and scores ≥66 indicate high vividness. In addition, the patient and a group of 18 neurotypical participants, aged 22-48, performed a computerized version of the *BIP - Battérie Imagerie-Perception* (Bourlon et al., 2009). The current version of the battery assesses imagery of object shapes (Fig. 1A), object colors (Fig. 1B), faces (Fig. 1C), letters (Fig. 1D) and spatial relationships on an imaginary map of France (Fig. 1E) (Bartolomeo, Bachoud-Lévi, Azouvi, & Chokron, 2005; Bourlon, Oliviero, Wattiez, Pouget, & Bartolomeo, 2011; Bourlon, Pradat-Diehl, Duret, Azouvi, & Bartolomeo, 2008). On each trial, participants hear a pair of nouns followed by an adjective: e.g. “cherries.… strawberries … dark”. They are requested to vividly imagine the items they hear. Immediately after, participants have to select the noun that is best represented by the adjective: in the previous example, it is “cherries”, which are darker than strawberries. Then they go on to indicate the overall vividness of their mental images on a 4-level Likert scale, pressing one of 4 buttons, where button 1 represents “no image at all” and button 4 represents a “vivid and realistic image”. RDS performed a slightly more extended version of the battery (18-20 items per imagery domain) than controls did (12 items per domain). A perceptual control task (Fig. 3F) employed the same stimuli used for the imagery tasks, except that stimuli were presented in an audio-visual format.

**Fig. 1.**
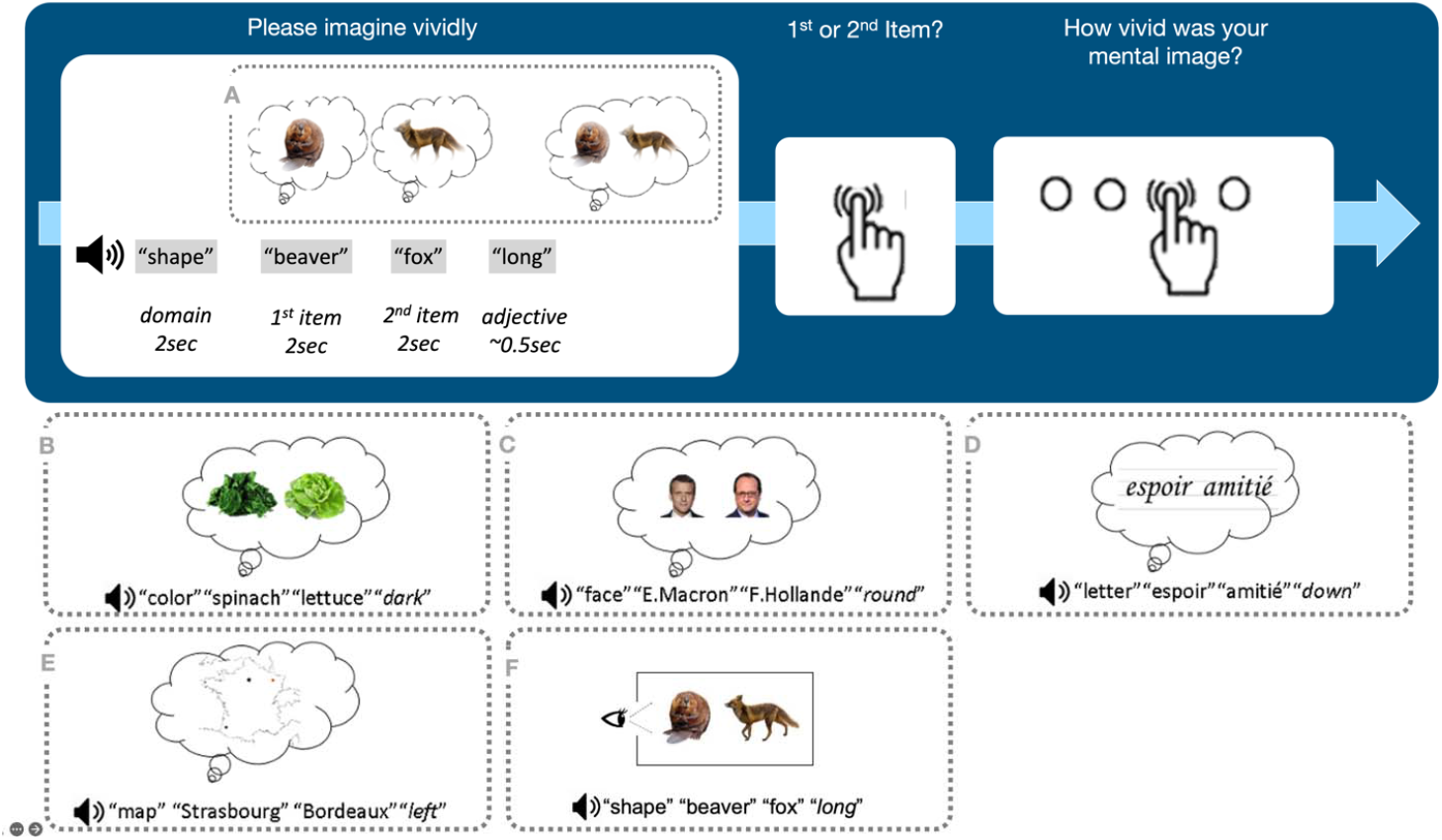
Examples of trials of the visual mental imagery battery performed by patient RDS. In different trials, participants are invited to decide about (A) the overall shape of animals or objects (round or long); (B) which fruit or vegetable is darker or lighter in color; (C) the general facial shape of celebrities (round or oval); (D) which handwritten word contains at least one ascender (t, l, d), or a descender (j, p, y); (E) which of 2 auditorily presented cities is right or left or Paris in an imaginary map of France. The perceptual task (F) is similar to the imagery tasks, except that stimuli are presented in an audio-visual format.

To compare RDS’s performance with controls’, we used SingleBayes_ES.exe, a tool for Bayesian analysis of the single case using a case-controls design (Crawford & Garthwaite, 2007). The program uses Bayesian Monte Carlo methods to test whether an individual’s score is significantly below the scores of controls such that the null hypothesis that it is an observation from the control population can be rejected, and provides a point estimate of the abnormality of the scores. Here, 2-sided p-values were considered as statistically significant at p < .05, and credible intervals are reported.

To assess the spatial relationships of the FIN with the patient’s lesion, we took the FIN volume from our meta-analysis (Spagna et al., 2021) and co-registered it to the patient’s T1 images in the native space.

To identify the fibers that are connected to the FIN, the functional ROI derived from our meta-analysis (Spagna et al., 2021) was used. We visualized the fibers of two major white matter tracts passing through the FIN volume: the occipito-temporal ILF (Catani, Jones, Donato, & ffytche, 2003), and the AF (Catani, Jones, & ffytche, 2005). Fiber tracking was done on the group average template constructed from a total of 1065 subjects (Fig. 3A), that comes with DSIstudio (https://dsi-studio.labsolver.org/). A multishell diffusion scheme was used, and the b-values were 990,1985 and 2980 s/mm^2^. The number of diffusion sampling directions were 90, 90, and 90, respectively. The in-plane resolution was 1.25 mm. The slice thickness was 1.25 mm. The diffusion weighted images were resampled at 2.0 mm isotropic. The b-table was checked by an automatic quality control routine to ensure its accuracy (Schilling et al., 2019). The diffusion data were reconstructed in the MNI space using q-space diffeomorphic reconstruction (Yeh & Tseng, 2011) to obtain the spin distribution function (Yeh, Wedeen, & Tseng, 2010). A diffusion sampling length ratio of 1.7 was used. The output resolution of is 2 mm isotropic. The restricted diffusion was quantified using restricted diffusion imaging (Yeh, Liu, Hitchens, & Wu, 2017). A deterministic fiber tracking algorithm (Yeh, Verstynen, Wang, Fernández-Miranda, & Tseng, 2013) was used with augmented tracking strategies (Yeh, 2020) to improve reproducibility.

To identify the fibers that are connected to the FIN, the functional ROI derived from a meta-analysis including 27 articles was used (Spagna et al., 2021) (specification of FIN: MNI coordinates −42, −54, 18). In order to grow this region into the white matter, a dilatation by 1 mm was applied in DSIstudio.

For fiber tracking, the anatomy prior of a tractography atlas (Yeh et al., 2018) was used to separately map “Inferior_Longitudinal_Fasciculus_L” and the “Arcuate_Fasciculus_L” with a distance tolerance of 16 mm. A seeding region was placed at “Inferior_Longitudinal_Fasciculus_L” and the FIN served as a ROI (Figure 3 in orange; Supplementary Video 1). The same procedure was also used to identify fibers of the the “Arcuate_Fasciculus_L” that are connected to the FIN (Figure 3 in blue; Supplementary Video 1). The anisotropy threshold, the angular threshold (between 15 to 90 degrees) and the step size (from 0.5 voxel to 1.5 voxels) were randomly selected. Tracks with length shorter than 10mm or longer than 200 mm were discarded. For visualization purposes, a total of 500 tracts were calculated. Topology-informed pruning (Yeh et al., 2019) was applied to the tractography with 16 iterations to remove false connections.

To visualize the spatial relationships of the brain lesion with the FIN connections, the patient’s structural MRI image was normalized into MNI space by using the normalization tool in the Brain voyager software (https://www.brainvoyager.com/). The patient’s brain lesion was then manually delineated by an experienced rater, by using the MRIcron software (https://www.nitrc.org/projects/mricron/).

For fiber tracking in the patient’s native space (Figure 3B), a DTI diffusion scheme was used, and a total of 60 diffusion sampling directions were acquired. The b-value was 1505 s/mm^2^. The in-plane resolution was 2 mm. The slice thickness was 2 mm. The diffusion MRI data were rotated to align with the AC-PC line. The tensor metrics were calculated using DWI with b-value lower than 1750 s/mm^2^ For fiber tracking the same procedure explained above was used. To match the patient’s anatomy, the functional ROI of the FIN (Spagna et al., 2021) was coregistered to the patient’s individual T1-weighted MRI using spm12 toolbox in Matlab. The resulting neuroanatomical localization in the patient space was checked and corrected by an experienced rater.

To identify the cortical areas that connected by the fibers passing through the FIN, as well as the the fibers that were damaged by the lesion, we used the connectivity matrix function in DSI studio. For the HCP1065 template, several matrices representing the number of fibers terminating within regions of the FreeSurferDKT_Cortical atlas (Desikan et al., 2006) were calculated. The connectivity matrix was calculated by using the count of the connecting tracks restricted to the left hemisphere. In creating the connectivity matrix the ROIs were specified as pass regions (Ghulam-Jelani et al., 2021). In total, 4 connectivity matrices were generated; two fascicles (AF or ILF) passing through two ROIs (FIN or Lesion).

## Results

RDS reported vivid visual mental imagery overall. His vividness score was 77, close to the ceiling score of 80 of the French version (Santarpia et al., 2008) of the VVIQ, and higher than the 66 cutoff score that Santarpia et al indicated for high vividness. RDS is thus a high vividness subject on the VVIQ. For each domain of the battery, the mean trial-by-trial subjective vividness was 3.8/4 (object shape) 2.7/4 (object color), 3.7/4 (letters), 3.1/4 (faces), and 3.5/4 (map of France). RDS’s performance on the battery (Table 1) revealed reasonably preserved abilities in all the tested imagery domains. No significant difference emerged between RDS’s performance and controls’, except for the map of France, where he was 72% correct. This level of performance was worse than controls’ (who were 75%-100% correct), but better than the 50% chance level (binomial test, *p* = .049).

**Table 1.** Performance (number and percentage of correct responses) of RDS and controls on the visual mental imagery battery

| Task | Control sample | | | Case's score | Significance test<sup>a</sup> | | Estimated percentage of the control population obtaining a lower score than the case<sup>b</sup> | | Estimated effect size ($z_{cc}$) | |
| --- | --- | --- | --- | --- | --- | --- | --- | --- | --- | --- |
|  | n | Mean | SD |  | t | p | Point | (95% CI) | Point | (95% CI) |
| Object Shape | 18 | 10.18/12<br>(85%) | 12.25% | 15/18<br>(83%) | -0.159 | .876 | 43.764% | 26.50% to 61.96% | -0.163 | -0.625 to 0.304 |
| Object Color | 18 | 9.19/12<br>(77%) | 16.45% | 16/20<br>(80%) | 0.178 | .861 | 56.944% | 38.74% to 74.09% | 0.182 | -0.286 to 0.646 |
| Letters | 18 | 11.62/12<br>(97%) | 5.16% | 19/20<br>(95%) | -0.377 | .711 | 35.540% | 19.50% to 53.89% | -0.388 | -0.861 to 0.098 |
| Faces | 18 | 10.13/12<br>(83%) | 9.07% | 14/20<br>(70%) | -1.395 | .181 | 9.052% | 1.85% to 22.44% | -1.433 | -2.087 to -0.758 |
| Map of France | 18 | 10.87/12<br>(91%) | 5.99% | 13/18<br>(72%) | -3.087 | .007 | 0.334% | 0.00% to 2.22% | -3.172 | -4.315 to -2.011 |

<sup>a</sup> Crawford & Howell (1998); the results are for a two-tailed test
<sup>b</sup> Crawford & Garthwaite (2002)

On the perceptual control task, RDS’s performance was at or near ceiling (range 95%-100% correct).

T1-weighted MRI (Fig. 2) showed that RDS’s ischemic lesion encompassed the calcarine sulcus, the lingual, fusiform, and parahippocampal gyri in the left hemisphere, as well as the callosal splenium. In the fusiform gyrus, the lesion’s lateral border corresponded to the midfusiform sulcus (see Fig. 2). Thus, the FG4 field, which lies lateral to the midfusiform sulcus (Lorenz et al., 2015) and contains the FIN (Spagna et al., 2021), was entirely spared by the lesion.

**Fig. 2.**
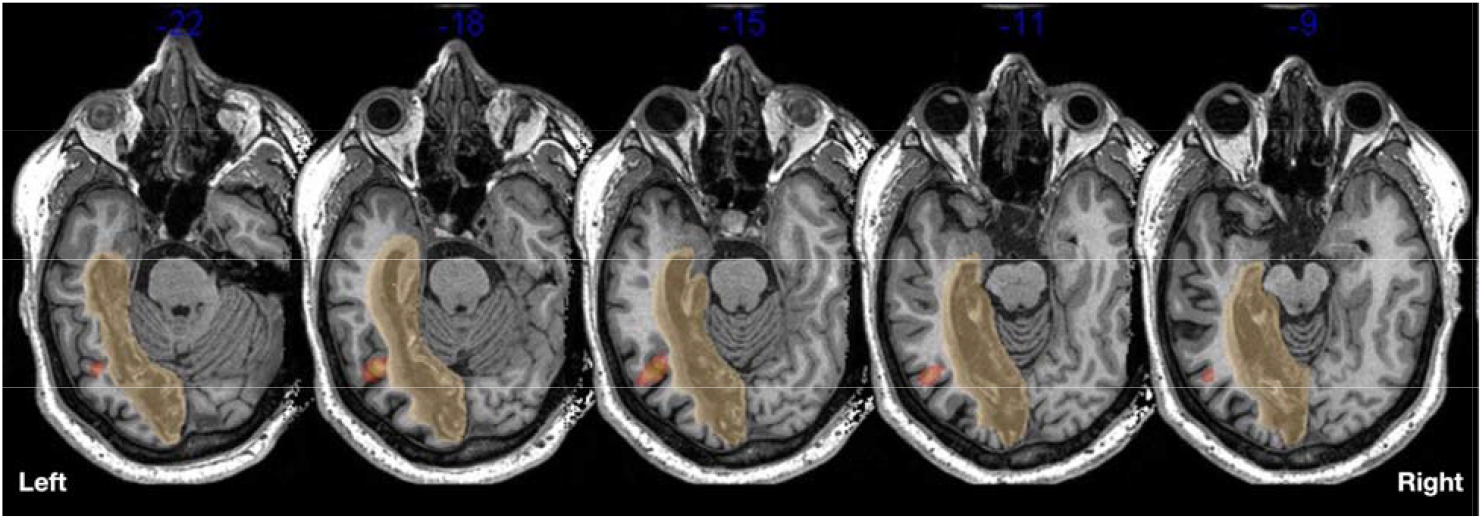
T1-weighted MRI showing RDS’s lesion (yellow) in native space, as well as the FIN location (orange).

White matter tractography (Fig. 3, supplementary video) demonstrated intact fibers passing through the FIN and belonging to two main systems: the ILF, linking the FIN to occipital regions and to anterior temporal regions important for semantic knowledge (Lambon Ralph et al., 2017), and the AF, connecting the FIN to perisylvian language circuits (Catani et al., 2005). Most intact ILF fibers connected the regions labeled as left_fusiform and left_inferior_temporal in the Desikan et al’s atlas (Desikan et al., 2006). Many ILF fibers leading to left_lateral_occipital, left_lingual, left_pericalcarine, left_superior_temporal were instead disconnected by the lesion. Concerning the AF, most intact fibers connected left_inferior_temporal to left_pars_opercularis, left_caudal_middle_frontal, and left_rostral_middle_frontal; fewer fibers connected left_inferior_temporal with left_pars_triangularis and left_precentral. The lesion mainly disconnected fibers going from left_inferior_temporal to left_caudal_middle_frontal.

**Fig. 3.**
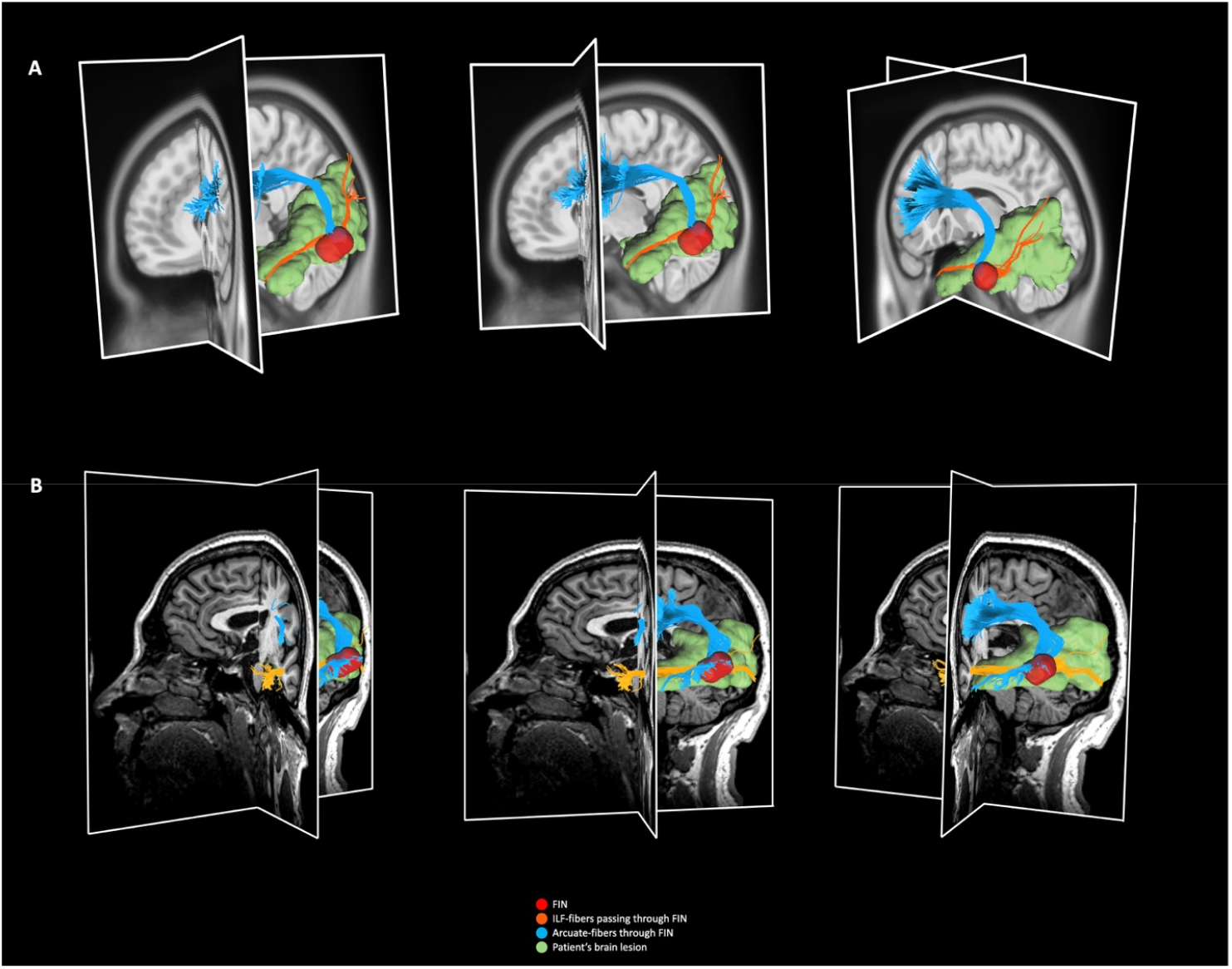
Connectional anatomy of the FIN (red), and its spatial relationship with a reconstruction of the patient’s lesion (green). Fibers passing through the FIN are visualized from two major tracts: the ILF (orange), which connects the FIN to occipital and temporal areas, and the AF (blue). Panel A shows the fibertracking on the HCP1065.2mm template, Panel B shows the fibertracking on the patient’s individual DTI, both with automated fiber tracking that come with DSIstudio (Version 2021.12.03 by Yeh; http://dsi-studio.labsolver.org/). The supplementary video https://bit.ly/3JrFYse) shows a 3D visualization of the ILF fibers and the AF fibers passing through the FIN, and their spatial relationship with the patient’s brain lesion.

## Discussion

The occipito-temporal stroke suffered by RDS extensively damaged the portion of the left fusiform gyrus situated medially to the midfusiform sulcus, but spared its lateral part, which contains the FIN (Spagna et al., 2021) in cytoarchitectonic sector FG4 (Lorenz et al., 2015). Lesion reconstruction and white matter tractography strongly suggest that the connectivity of the FIN with the anterior temporal lobe and with language circuits was also spared by the lesion. RDS’s intact introspection and reasonably preserved performance for visual mental imagery, together with the preservation of the FIN, is thus consistent with the hypothesis of a causal implication of the FIN in this ability. Moreover, in this patient the FIN could not communicate with the left primary visual cortex, which was totally destroyed by the lesion; as a consequence, our data support the additional hypothesis that visual mental imagery engages the FIN in a topdown fashion, perhaps through the ILF from the anterior temporal lobe important for semantic processing (Lambon Ralph et al., 2017), and through the AF from prefrontal regions and language circuits. The present results thus add to extensive neuropsychological evidence (Bartolomeo et al., 2020; Liu et al., 2021; Spagna, 2022) against the dominant model of visual mental imagery, which emphasizes the role of primary visual cortex in this ability (Kosslyn et al., 1999; Kosslyn, Thompson, Kim, & Alpert, 1995; Pearson, 2019). In principle, however, other spared occipito-temporal regions could contribute to RDS’s visual mental imagery. For example, primary visual cortex was intact in the right hemisphere. However, it is of note that, in the present patient, splenial disconnection (which was demostrated in a previous study using white matter tractography, see Siuda-Krzywicka et al., 2019) prevented any direct communication between the right, intact primary visual cortex and the posterior part of the left hemisphere. Thus, putative contributions of the right hemisphere primary visual cortex to FIN activity should have traveled indirectly through more anterior callosal connections.

Our conclusions are based on RDS’s performance seven years after his stroke. This extended time lapse could have allowed plastic phenomena to occur. Against this possibility, we note that RDS’s deficits in reading and color naming were stable over the years. Moreover, testing at closer temporal intervals from the stroke may suffer from another potential confounding factor, the occurrence of diaschisis phenomena (Bartolomeo, 2011). For example, in patients with severe strokes, gray matter blood flow in the homolog regions of the opposite hemisphere can be decreased for several months, and may approach normal levels only 12-24 months post-stroke (Meyer, Obara, & Muramatsu, 1993). Note, however, that even seven years post-stroke RDS’s face imagery scores were numerically inferior to those for other domains. Face imagery may engage the right-hemisphere fusiform face area (O’Craven & Kanwisher, 2000), and thus require interhemispheric integration. In RDS, perhaps interhemispheric communication was still partially dysfunctional at the time of testing, as a consequence of splenial disconnection. A similar point might be advanced for RDS’s impaired performance on the test based on the map of France, which is also likely to require interhemispheric integration (Rode et al., 2010). Further evidence from the follow-up of brain-damaged patients, coupled with advanced neuroimaging, is needed to discriminate between the sometimes contradictory roles of the nondamaged hemisphere in post-stroke cognitive deficits (Bartolomeo & Thiebaut de Schotten, 2016).

The present evidence only indirectly speaks to the causal role of the FIN in visual mental imagery. However, our results nicely complement more direct evidence coming from brain-damaged patients with impaired visual mental imagery. For example, two patients found themselves unable to build visual mental images after a closed head trauma (Moro, Berlucchi, Lerch, Tomaiuolo, & Aglioti, 2008). In both cases, the damage affected the left BA 37, and was likely to include the lateral portion of the fusiform gyrus with the FIN. Traumatic brain injuries, like those suffered by Moro et al.’s patients, typically provoke diffuse axonal injury, which is likely to disrupt white matter connectivity in the large-scale brain networks supporting visual mental imagery (Bartolomeo, 2008; Mechelli, Price, Friston, & Ishai, 2004). Thus, FIN dysfunction leading to impaired visual mental imagery should not be interpreted in a localist way, but as a source of perturbation of large-scale brain networks (Bartolomeo, 2011). Another patient with left temporal damage had impaired perception and imagery for orthographic material, but not for other domains (Bartolomeo, Bachoud-Lévi, Chokron, & Degos, 2002). Damage or disconnection of domain-preferring regions such as the visual word form area (Dehaene & Cohen, 2011) may account for such domain-selective deficits of visual mental imagery, as opposed to domain-general imagery deficits which, in the present framework, may instead result from FIN dysfunction.

In apparent contradiction with the present evidence, a recent case report (Thorudottir et al., 2020) described patient PL518, an architect who spontaneously complained to have become unable to visualize items after a bilateral stroke in the territory of the posterior cerebral artery. The lesion included the left medial fusiform gyrus, but spared its lateral portion. However, the lesion did extend more laterally in the fusiform white matter than similar lesions in patients without visual mental imagery deficits (see their Fig. 3A). Thus, disconnections within the fusiform white matter might have contributed to FIN dysfunction and consequent visual mental imagery impairment in PL518, perhaps in combination with the extensive accompanying lesions in the right hemisphere, which might have deprived the left hemisphere of potential interhemispheric compensation (Bartolomeo & Thiebaut de Schotten, 2016).

Taken together with these previous studies, the present evidence suggests a crucial role for the FIN in visual mental imagery. Specifically, the FIN might integrate, on the one side, elements of semantic knowledge stored in the anterior temporal lobe (Lambon Ralph et al., 2017; Persichetti, Denning, Gotts, & Martin, 2021), and distributed linguistic representations (Popham et al., 2021), with, on the other side, high-level visual representations in domain-preferring regions in the ventral temporal cortex (Mahon & Caramazza, 2011). Following the lead of Heinrich Lissauer’s seminal ideas (Bartolomeo, 2021; Lissauer, 1890; Lissauer & Jackson, 1988), we propose that dissociations in performance between perceptual and imagery abilities may emerge when the FIN or other high-level visual regions in the ventral temporal cortex are deafferented from perceptual input processed in more posterior regions. Such posterior disconnections would result in impaired perception with preserved imagery (Bartolomeo et al., 1998). The present evidence suggests that, in these cases, visual mental imagery would be supported by the ILF and AF connections to the FIN.

A more direct test of this model would have required fMRI evidence of spared FIN activity during visual mental imagery in our patient; unfortunately, however, when we attempted such an experiment RDS had a panic reaction in the MRI machine, and subsequently declined to participate in any further neuroimaging exams. As a consequence, the localization of the FIN in the patient’s brain must be considered as a (likely) approximation, because it derives from our meta-analysis of fMRI studies (Spagna et al., 2021). Finally, we note that although single patient studies can achieve a level of detail unattainable in group studies, without extensive replication one cannot always exclude the influence of idiosyncratic variations of the mind/brain (Bartolomeo, Seidel Malkinson, & de Vito, 2017). This seems, however, unlikely for RDS, because there are reasons to consider his premorbid neurocognitive profile as representative of the general population (Siuda-Krzywicka et al., 2019). Accumulating evidence from in-depth studies of other patients, with brain lesions inducing or not deficits of visual mental imagery, will be important to confirm or refute the present model.

